# Bioinformatic Characterization of Regulated IRE1α-Dependent Decay (RIDD) in Heart Failure

**DOI:** 10.64898/2026.08.20.745896

**Authors:** Nisha Bhattarai, Angelina M. Kendi, Michael W. Stoner, Sruti S. Shiva, Brett A. Kaufman, Iain Scott

## Abstract

Inositol-requiring enzyme 1α (IRE1α) is a canonical signaling factor in the unfolded protein response (UPR). In addition to this essential role (which prevents the accumulation of misfolded proteins in the endoplasmic reticulum), the endoribonuclease activity of IRE1α targets multiple mRNAs for degradation through a process called Regulated IRE1α-Dependent Decay (RIDD). The products of over 50 genes have been identified as RIDD targets; however, the biological significance of this process remains underexplored. Using publicly available datasets, we examined the fate of 27 well-characterized RIDD targets in the septal wall of heart failure patients, and in mice subject to pressure overload-induced heart failure. We show that decreased mRNA abundance from these RIDD substrate genes – an outcome consistent with RIDD induction – is commonly observed in heart failure.

## Introduction

Inositol-requiring enzyme 1α (IRE1α) is the key regulatory enzyme in one of the three pathways of the unfolded protein response (UPR), which prevents the accumulation of misfolded proteins in the endoplasmic reticulum (ER; reviewed in Lemmer et al., 2021). When activated by ER stress or unfolded proteins in the ER lumen, the endoribonuclease domain of IRE1α cleaves the mRNA of X-box binding protein 1 (XBP1) to remove an intron, converting inactive XBP1 into XBP1s, its active transcription factor form. XBP1s then translocates from the ER to the nucleus, activating the transcription of chaperone proteins that help mitigate protein misfolding in the ER (Lemmer et al., 2021).

In addition to this canonical activity in the UPR, the endoribonuclease activity of IRE1α cleaves the mRNA of multiple genes as they transit the ER, in a process termed Regulated IRE1α-Dependent Decay (RIDD) (Hollien and Weissman, 2006; Hollien et al., 2009). During RIDD, IRE1α recognizes a semi-conserved stem-loop structure on mRNAs, which are then degraded via the enzyme’s endoribonuclease activity (Bright et al., 2015). Given that at least 50 potential substrate mRNAs have now been identified as being degraded by RIDD during ER stress (Bright et al., 2015), this process can be viewed as having a significant effect on cellular proteostasis. However, at the present time, our understanding of RIDD biology in the failing heart remains underdeveloped. To address this limitation, we examined the potential induction of RIDD in published databases of human and mouse heart failure samples and experimentally tested the induction of RIDD *in vitro*. Our findings suggest that RIDD is likely to be a commonly observed process in the failing heart.

## Methods

### Bibliographic analysis

A list of 27 RIDD targets, previously identified in at least two separate peer-reviewed publications, was obtained from Bright et al. (2015). Human heart failure or other cardiovascular disorder (CVD) publications that focused on these genes were obtained from Pubmed (https://pubmed.ncbi.nlm.nih.gov/) between 25-27^th^ July 2026 using the search terms “[GENE NAME] + Human + Heart Failure” or “[GENE NAME] + Human + Heart”, respectively. Following each search, a manual review process was used to ensure that human disease mechanisms for either heart failure or a specified cardiovascular disorder (e.g. hypertension, dilated cardiomyopathy, pulmonary hypertension, etc.) were investigated in at least one peer-reviewed publication for that specific gene.

### Human heart RNA-seq data

RNA-seq read data from control (n = 25), HFrEF (n = 33), and HFpEF (n = 32) patients were obtained from a dataset published by Hahn et al. (2021). Data for each of the 27 RIDD targets were manually extracted using Ensembl stable ID codes, and were allocated to groups (control, HFrEF, or HFpEF) using the supplied deidentified patient IDs. Mean ± SEM values for each group were calculated and subject to statistical analysis.

### Mouse heart RNA-seq data

RNA-seq read data from control (sham surgery) and heart failure (8 weeks after TAC surgery) mice (n = 3 per group) were obtained from a dataset published by Froese et al. (2022). Data for each of the 27 RIDD targets were manually extracted and subject to statistical analysis as above.

### Pathway enrichment analysis

Gene-based pathway enrichment analysis was performed using ShinyGO 0.85 (https://bioinformatics.sdstate.edu/go/). Pathway number was set to 10, and outputs were analyzed via the GO Biological Process database.

### Cell culture and treatments

Human HepG2 cells were cultured under standard conditions (25 mM glucose DMEM, 10% FBS, antibiotic-antimycotic, 5% CO_2_, at 37°C; all ThermoFisher, USA) and treated with 1 μg/mL tunicamycin or 0.1 αg/mL brefeldin A for 24 h.

### RT-qPCR and Western Blot

RNA was harvested from cells using the RNEasy kit (Qiagen), with a μLITE analyzer (BioDrop, USA) used to evaluate the quality and quantity. RNA (1 μg) was used to synthesize cDNA using the Maxima Reverse Transcriptase (ThermoFisher, USA) kit, then gene expression studies were performed with specific primers for *BLOC1S1* (Cat.No. QT00016002), and *GAPDH* (Cat.No. QT00079247; all Qiagen, USA).

For western blot analysis, cells were lysed in 1% CHAPS buffer. Protein samples were quantified using a μLITE analyzer (BioDrop), and 20 μg of proteins was separated on an SDS-PAGE gel.Protein bands were transferred onto 0.2/0.45 μm nitrocellulose membranes, which were blocked using Odyssey blocking buffer (Li-Cor, USA). Membranes were incubated with primary antibodies for GAPDH (Cell Signaling, Cat. 97166 [D4C6R], mouse mAb) or BLOC1S1 (Bugga et al., 2024; rabbit pAb)

### Statistics

Statistical analysis was performed using Graphpad Prism v.11. Data were analyzed using multiple t-tests with post-hoc adjustment for multiple comparisons.

## Results

### Bioinformatic analysis of RIDD in heart failure

Using a curated list of 27 recognized RIDD targets (Bright et al., 2015), we first examined whether these RIDD substrates have been associated with heart failure or other human cardiovascular diseases. Bibliographic analysis of the Pubmed database demonstrated that over one-third (37%) of the RIDD substrates examined have been implicated in heart failure previously (**Figure 1A**). When including cardiovascular diseases of any etiology, two-thirds (67%) of the RIDD targets examined have been linked to human heart disease (**Figure 1A**). These data suggest that RIDD induction may be a common feature of cardiac disease pathophysiology. We next examined what biological pathways are highly enriched between the different RIDD substrates and found that both isoprenoid and lipid metabolism are overrepresented in the 27 genes (**Figure 1B**). We next sought to determine how different heart failure etiologies may be associated with RIDD induction using a published human dataset (Hahn et al., 2021). A majority (14 of 27) of RIDD substrate genes in patients with heart failure with reduced ejection fraction (HFrEF) displayed reduced mRNA abundance, while 12 of 27 of RIDD substrates showed reduced mRNA abundance in patients with heart failure with preserved ejection fraction (HFpEF) (**Figure 1C**). About half of the reduced mRNA abundance observed in RIDD substrate genes was shared between the two heart failure etiologies, with five genes being specific to HFrEF (*NDUFA9, PMVK, NRM, HGSNAT*, and *BLOC1S1*), and three being specific to HFpEF (*TSPAN3, TPP1*, and *PDAP1*) (**Figure 1D**).

**Figure 1.**
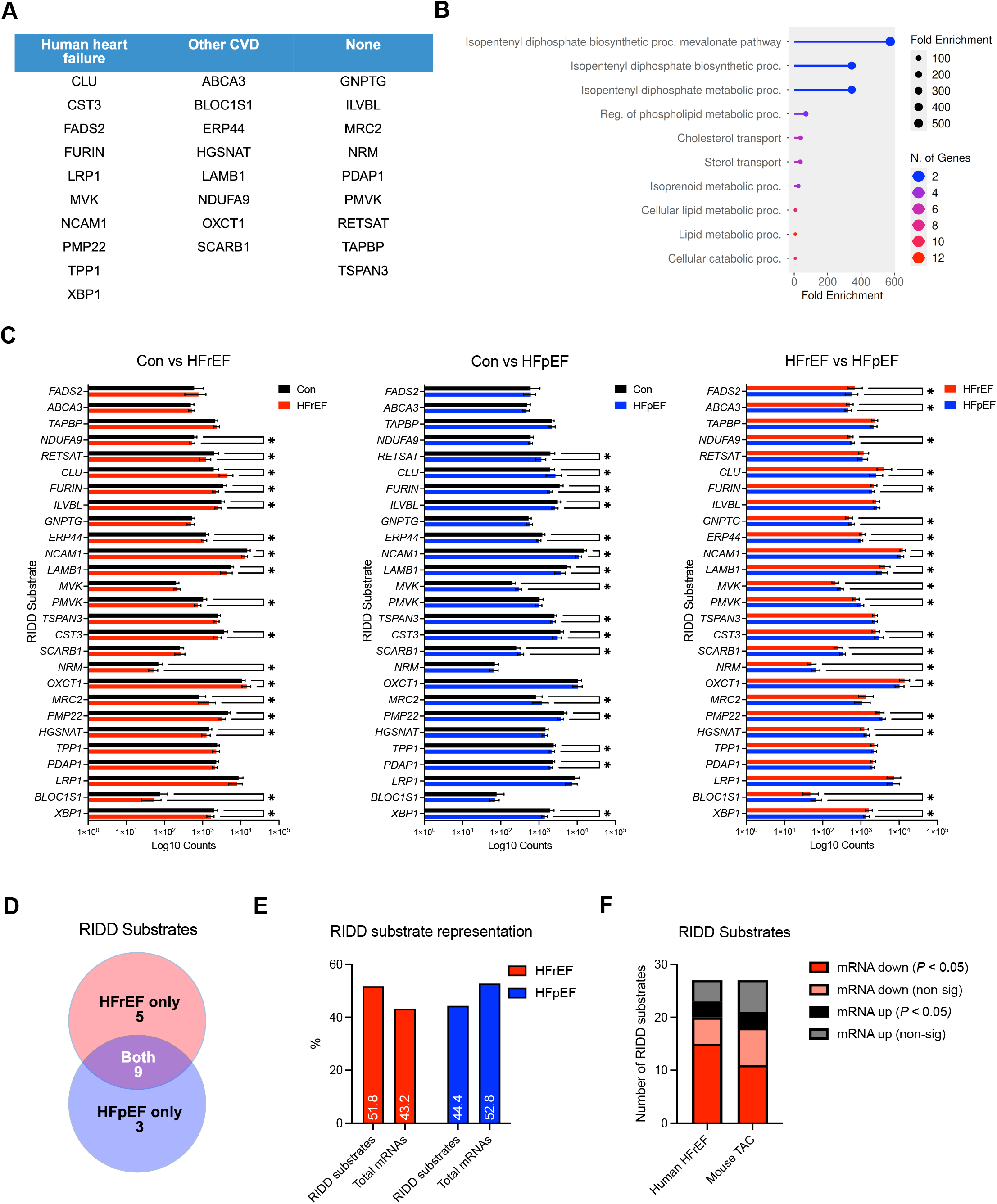
Bioinformatic analysis of RIDD induction in hearts from HFrEF and HFpEF patients. **A**. Bibliographic searches demonstrate that a majority of known RIDD substrates have been implicated in heart failure or other cardiovascular diseases (CVD). **B**. Gene enrichment analyses of known RIDD substrates suggest that pathways related to isoprenoid and lipid metabolism are highly represented. **C**. Analysis of mRNA abundance of known RIDD substrates in healthy control, HFrEF, and HFpEF patient samples. Data are mean ± SEM, * = *P* < 0.05. **D**. Number of RIDD substrates with significantly decreased mRNA abundance in either HFrEF, HFpEF, or shared between both heart failure etiologies. **E**. Significantly decreased RIDD substrate mRNAs are more common in HFrEF samples when compared to significant decreased in the total transcript population. In contrast, decreased RIDD substrate mRNAs are less common when compared to significant decreases in the total population of transcripts in HFpEF samples. **F**. Comparison of RIDD substrate mRNA levels in human HFrEF and mouse pressure overload-induced heart failure (TAC) suggests a similar level of RIDD induction.

We next examined whether the reduced mRNA expression observed in RIDD genes matched the general mRNA population (i.e., would suggest a non-specific effect that was common to both RIDD and non-RIDD substrates), or whether it was found at a higher rate in RIDD genes (i.e., would suggest an effect specific to RIDD substrates). In HFrEF, 51.8% of the RIDD substrate genes showed a significant decrease in mRNA abundance, relative to 43.2% of the total mRNA population (**Figure 1E**). These data suggest that RIDD genes are targeted for mRNA reductions at a higher rate than the transcriptome as a whole, implying that RIDD may be specifically upregulated in HFrEF. In contrast, the proportion of the total transcriptome that is significantly decreased in HFpEF is higher (52.8%) than the number of significantly decreased RIDD substrate genes under the same condition (44.4%) (**Figure 1E**). This suggests that RIDD induction is lower in HFpEF than in HFrEF, and that the RIDD program may be a more important feature of the latter disease.

We next examine whether RIDD induction was a consistent finding across different HFrEF conditions. Analysis of an RNA-seq database from a mouse model of pressure overload-induced heart failure (Froese et al., 2022) demonstrated that a similar proportion of RIDD substrate gene mRNAs were reduced in mouse and human HFrEF (**Figure 1F**). These data suggest that RIDD induction may be a common feature of heart failure with reduced ejection fraction, regardless of species or etiology.

### Confirmation of BLOC1S1 as a RIDD target in human cells

Finally, we confirmed whether the induction of RIDD would lead to the reduced abundance of a key RIDD target, *BLOC1S1* (Bright et al., 2015). Human HepG2 cells treated with ER stress-inducing chemicals tunicamycin or brefeldin A for 24 hour displayed a significant decrease in mRNA abundance consistent with RIDD induction (**Figure 2A**). Consistent with this finding, there was a significant decrease in BLOC1S1 protein abundance in brefeldin A treated cells, along with a decrease in protein in tunicamycin treated cells that trended towards significance (*P =* 0.057) (**Figure 2B**). Combined, these data support the finding that BLOC1S1 is a RIDD target in human cells, and induction of this program affects both its mRNA and protein abundance.

**Figure 2.**
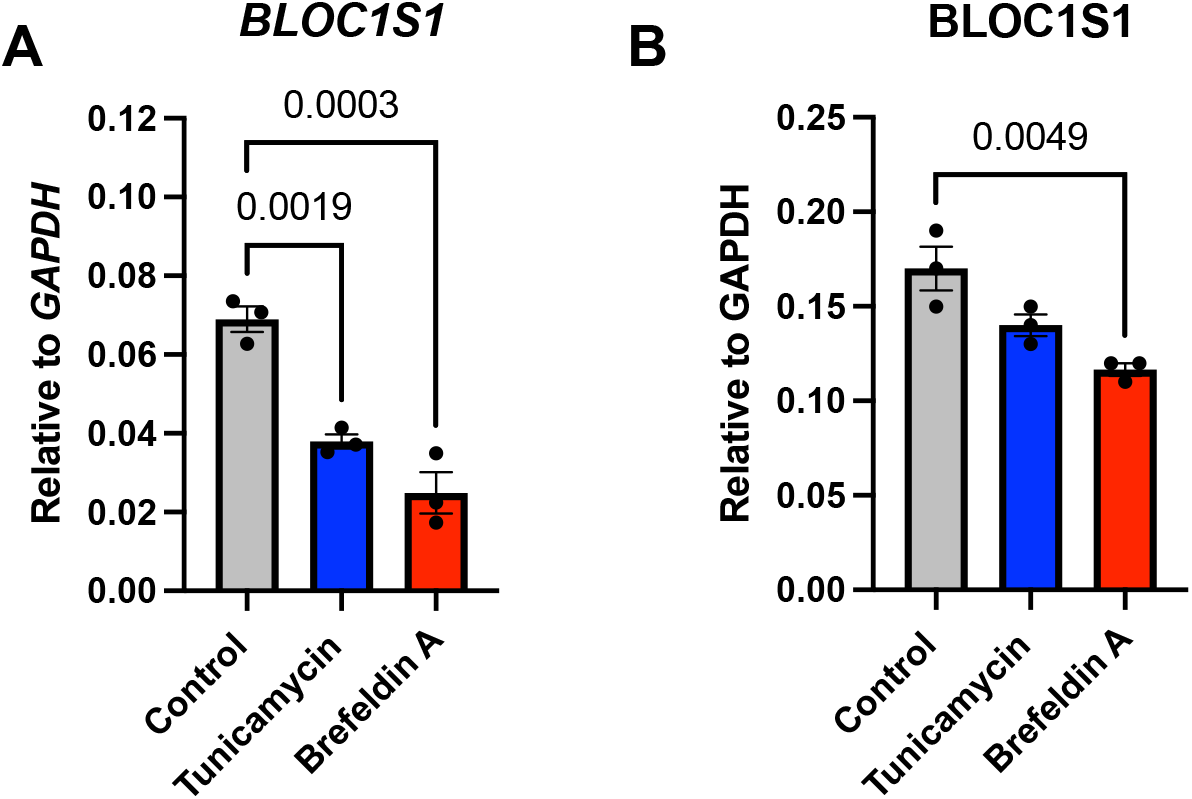
Experimental confirmation that induction of RIDD via ER stress leads to downregulation of BLOC1S1. **A**. mRNA abundance of *BLOC1S1* in human HepG2 cells after treatment with Tunicamycin or Brefeldin A for 24 h. **B**. Protein abundance of BLOC1S1 in human HepG2 cells after treatment with Tunicamycin or Brefeldin A for 24 h.

## Discussion

Using transcriptomic analysis, we show that over 60% of well-characterized RIDD substrates display decreased mRNA expression in all types of heart failure, consistent with the induction of the RIDD program. While ER stress and the UPR are central to the development of various cardiac disorders, the regulatory role played by RIDD is not well understood. It has been shown that the mRNA of Corin – a serine protease that processes natriuretic peptides like pro-ANP and pro-BNP into their active forms – is degraded by RIDD during the progression of heart failure, which may negatively impact overall control of natriuretic peptide system (Lee et al., 2015). The same authors also demonstrated that over 200 mRNAs were decreased greater than two-fold in end-stage heart failure samples compared to healthy hearts, and that the majority of these were ER-transiting mRNAs that may possibly be subject to IRE1α-depdendent processing (Lee et al., 2015).

Interestingly, the level of RIDD induction in HFrEF appears to be higher than in HFpEF, suggesting that diverse heart failure etiologies may result in a differential induction of this program in patients (**Figure 1E**). This finding mirrors the differences observed in XBP1 processing by IRE1α between mouse models of HFpEF (high fat diet + L-NAME) and HFrEF (severe transaortic constriction), where XBP1s levels were significantly decreased in HFpEF but unchanged in HFrEF (Schiattarella et al., 2019). As such, the etiology of heart failure may have a strong regulatory effect on the induction of IRE1α-dependent decay. Further study of the RIDD process in the heart, along with its effect on specific substrates, may yield novel therapeutic avenues to tackle heart failure.

## Funding

This work was supported by National Institute of Health Research Grants (R01HL147861, R0HL156874) and American Heart Association Established Investigator Award (23EIA1037834) to I.S.

## Author contributions

N.B., A.M.K., and M.W.S executed bioinformatic and experimental procedures, analyzed data, and edited the manuscript. S.S.S. and B.A.K. provided critical expertise. I.S. planned bioinformatic analyses, performed secondary analysis of data, generated figures, and wrote the manuscript. All authors have reviewed the final manuscript and agree with the submission.

## Competing interests

The authors declare that they have no competing financial or ethical interests related to the submission and publication of this article.

## Notes

### Competing Interest Statement

The authors have declared no competing interest.

